# Comparison of reprogramming factor targets reveals both species-specific and conserved mechanisms in early iPS cells

**DOI:** 10.1101/225326

**Authors:** Kai Fu, Constantinos Chronis, Abdenour Soufi, Giancarlo Bonora, Miguel Edwards, Steve Smale, Kenneth S. Zaret, Kathrin Plath, Matteo Pellegrini

## Abstract

Both human and mouse fibroblasts can be reprogrammed to pluripotency with Oct4, Sox2, Klf4, and c-Myc (OSKM) transcription factors. While both systems generate pluripotency, human reprogramming takes considerably longer than mouse. To assess additional similarities and differences, we sought to compare the binding of the reprogramming factors between the two systems. In human fibroblasts, the OSK factors initially target many more closed chromatin sites compared to mouse. Despite this difference, the intra- and intergenic distribution of target sites, target genes, primary binding motifs, and combinatorial binding patterns between the reprogramming factors are largely shared. However, while many OSKM binding events in early mouse cell reprogramming occur in syntenic regions, only a limited number is conserved in human. In summary, these findings suggest similar general effects of OSKM binding across these two species, even though the detailed regulatory networks have diverged significantly.

## Introduction

By expressing the transcription factors Oct4, Sox2, Klf4 and c-Myc (abbreviated as OSKM), differentiated cells can be reprogrammed into induced pluripotent stem cells (iPSCs) that have the ability to differentiate into any type of cell (Takahashi & Yamanaka, 2006a; Takahashi et al., 2007). iPSC technology holds great promise in regenerative medicine and for the modeling of diseases in a culture dish (Hirschi, Li, & Roy, 2014; Singh, Kalsan, Kumar, Saini, & Chandra, 2015). However, there is still limited understanding of the essential mechanisms underlying reprogramming of somatic cells to iPSCs. Furthermore, there are marked differences in the reprogramming process for mouse and human cells, even though reprogramming can be accomplished by the same set of factors. Mouse cells reprogram within a week or two, whereas human cells take up to a month and the efficiency of the conversion is typically lower in the human system (Takahashi & Yamanaka, 2016; Yamanaka, 2012). Moreover, while mouse cells can be reprogrammed efficiently with OSK alone, ectopic c-Myc expression is more critical in the human process (Malik & Rao, 2013; Takahashi et al., 2007; Takahashi & Yamanaka, 2006b). To understand universal features of reprogramming across species, we characterized the differences and similarities in the regulatory networks that were manifested at the onset of reprogramming of human and mouse somatic cells.

An important approach towards understanding the reprogramming process is to systematically investigate the binding of reprogramming factors in the genome. By investigating OSKM binding at 48 hours of reprogramming, previous studies have begun to elucidate the patterns and regulatory roles of OSKM in early reprogramming in the human and mouse systems (Chronis et al., 2017; Soufi, Donahue, & Zaret, 2012; Soufi et al., 2015). Reprogramming typically is an inefficient process where only few cells in the culture dish induce the pluripotency program, yielding a highly heterogeneous cell population at the end of the process. However, in the first 48 hours of reprogramming, the reprogramming culture is thought to react homogeneously (Koche et al., 2011; Polo et al., 2012), enabling location studies of OSKM in the early reprogramming population. Moreover, for the 48-hour time point in mouse, we used fetal bovine serum containing media, which results in iPSC colonies within 2-3 weeks. In these conditions, the timing of reprogramming is similar to that found in human experiments. The early human and mouse cells are thus expected to be in a similar stage of reprogramming. However, the final iPSC stage between human and mouse is significantly different: the human cells are reprogrammed to a primed stage while the mouse cells are reprogrammed to a naïve stage (Nichols & Smith, 2009). For this reason, in this study we focused on the 48-hour comparison instead of the iPSC stage of reprogramming.

In this study, we compared the initial OSKM binding events between human and mouse fibroblasts to shed light on both conserved and species-specific mechanisms of OSKM-mediated processes early in reprogramming. By focusing on the binding events of OSKM early in reprogramming, we guaranteed minimal influence of the differences between human and mouse cell reprogramming that resulted in mouse iPSCs in the naïve pluripotent state and human iPSCs in the primed pluripotent state caused by the external culture conditions. We first show that general features of OSKM binding events, such as inter- and intragenic distribution, target genes, primary binding motifs, and combinatorial binding patterns between the reprogramming factors, are largely similar between human and mouse. However, when we compared the locations of OSKM binding events, we found that only a small fraction of binding sites in syntenic regions were conserved between human and mouse at 48 hours of reprogramming. This result indicates that the binding of the reprogramming factors is in large part distinct at the initial stage of the reprogramming process. We show that conserved binding events within syntenic regions often represent target sites that are also bound in the pluripotent end state and tend to occur in promoters and enhancers, suggesting that the engagement of pluripotency sites early in reprogramming is a conserved mechanism between mouse and human reprogramming. Lastly, we show that both motif usage and chromatin states contribute to the conservation of binding events in early human and mouse reprogramming.

## Results

### General features of OSKM binding events in early human and mouse reprogramming

In this study, we compare the binding of OSKM peaks in mouse and human at 48 hours post transfection. This is accomplished by analyzing previously published datasets (Chronis et al., 2017; Soufi et al., 2012). We note that there are some differences in the mouse and human datasets that are due to the difference in overexpression methodology and the starting cell type. While the mouse data was generated by overexpressing the pluripotency factors using a polycistronic cassette (ensuring that each cell expresses all four factors at comparable levels), the human data was generated using individual lentiviral vectors, which leads to more variability in the combination and level of expression of the factors. However, as we show below, these differences do not have a significant impact on our conclusions.

We first addressed the effects of overexpression between polycistronic and individual based approaches. While the primary results presented in Chronis *et al* were based on a polycistronic cassette (Chronis et al., 2017), in the same study we also collected binding data generated by individually overexpressing factors using pmX. We showed that in mouse, individual retroviral based expression of Oct4, Sox2 and Klf4 (OSK) have strong signals in the polycistronic derived OSK peaks, indicating that the OSK signal from the two systems are enriched in similar genomic loci (Supp Fig 1). Moreover, we note that while the mouse experiments were carried out in embryonic fibroblasts, the human studies were done in fetal foreskin fibroblast. Since we did not have access to epigenomes from both embryonic fibroblasts and fetal foreskin fibroblasts in either human or mouse, we were not able to compare the potential differences between the two starting cell types. Nonetheless, we do have access to epigenomes for both human foreskin newborn and human lung fetal fibroblasts from Roadmap Epigenomics Project. To address the potential differences between different types of fibroblasts, we used DNaseI hypersensitive sites to represent the chromatin states and then compared their overlapping. Supplementary Fig 2 then shows that the two types of fibroblasts have a large overlapped number of DNaseI peaks, suggesting the overall similarities of chromatin states between those two types of fibroblasts. We thus argue that chromatin changes are modest between the two types of fibroblasts we used in this study.

Having shown that these two experimental strategies yield similar OSKM binding events, we chose to focus our analyses on the mouse polycistronic and human individual lentiviral cassette where all our ChIP-Seq and RNA-Seq data was collected. To further enable their comparison, we generated OSKM peaks for both human and mouse cells reprogramming using the same analysis pipeline for mapping and peak calling, setting the peak calling q-value cutoff of 0.05 (see Methods).

The human and mouse data sets generated a similar number of peaks for Oct4, while the early human reprogramming culture had about twice as many peaks for the other three factors compared to the mouse (Supp Fig 3). In both human and mouse, ChIP-seq for Myc generated fewer peaks than O, S, or K (Supp Fig 3). The average fragment size (average distances between plus strand reads and minus strand reads) was similar for all four reprogramming factors in the mouse and human data sets (Supp Fig 4, Supp Fig 5). We found that the human datasets for O, S, and K had a lower signal to noise ratio than the mouse data sets, whereas M binding events were slightly stronger in the human sample (Supp Fig 4, Supp Fig 5, Supp Fig 6).

We first asked whether OSKM peaks had a similar positional distribution with respect to transcriptional start sites (TSSs) in the two species (Fig 1a). Specifically, we classified the distances between peaks and TSSs into different groups, i.e. 0 to 5kb, 5 to 50kb etc. We found that O, S and K peaks were most abundant in the −500 to −50kb and 50 to 500kb bins in both human and mouse, indicating that O, S and K in both human and mouse predominantly bind regions distal to TSS. M peaks, however, were most abundant in −500 to −50kb and 50 to 500kb bins in human, while most abundant in the −5 to 0kb and 0 to 5kb bins in mouse. This result reveals that M has a different distribution between human and mouse: in humans M tends to bind distally to the TSS whereas in mouse it tends to bind proximal to TSS regions. In addition, we observed less overall binding of O, S and K in proximity to the TSS compared to distal sequences in human than in mouse cells (Fig 1a).

**Figure 1.**
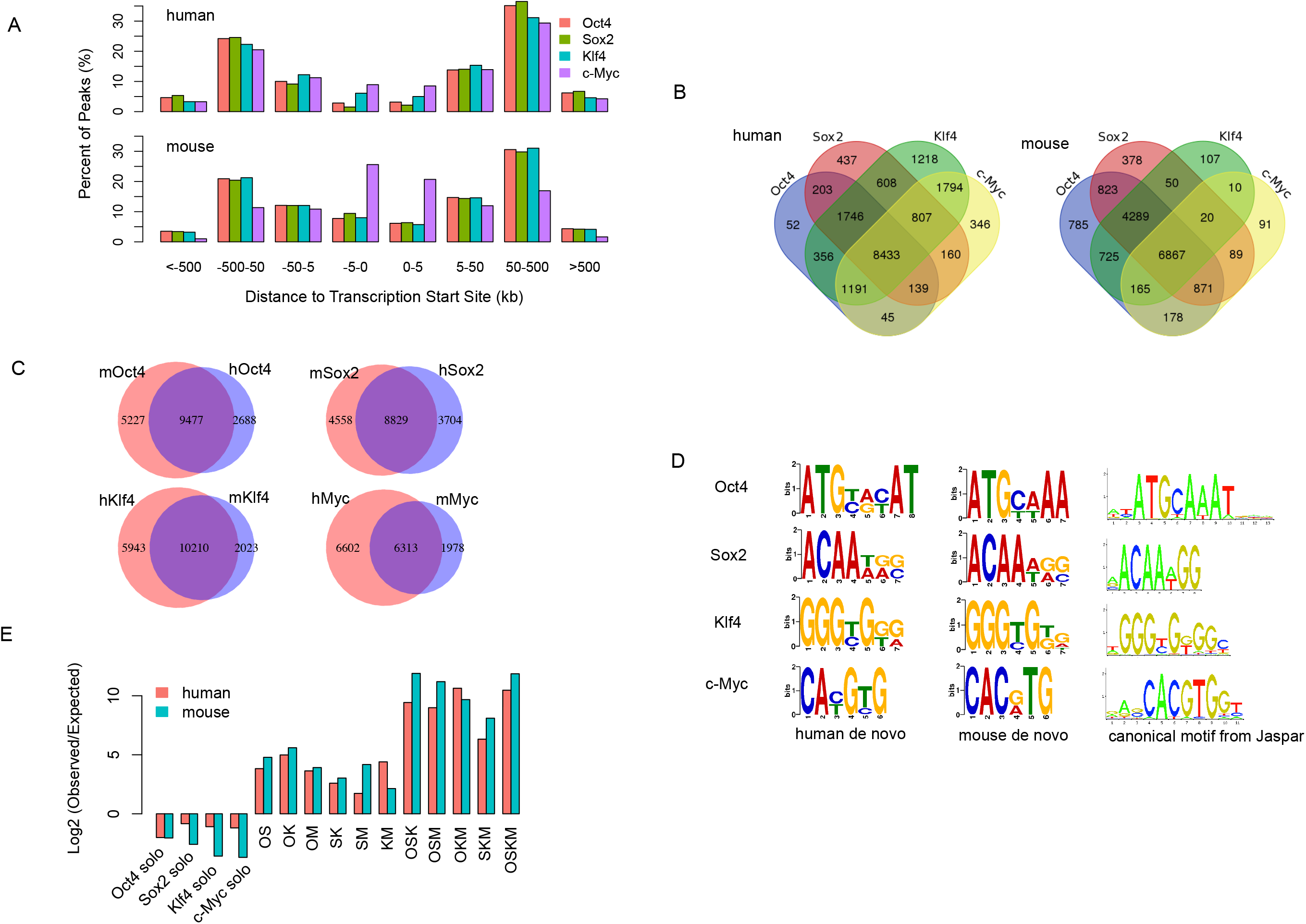
General feature comparison of OSKM ChIP-Seq peaks between human and mouse 48 hours fibroblast reprogramming. A. Positional distribution of OSKM peaks with respect to Transcription Start Sites (TSSs). The top panel shows the peaks in human while the bottom panel shows that in mouse. B. Venn diagram of OSKM co-targeted genes in human (left panel) and in mouse (right panel). C. Venn diagram of OSKM co-targeted orthologous genes between mouse and human. D. De novo and canonical motifs of OSKM peaks. E. Log2 ratio of observed combinatorial binding events versus expected.

We next compared the target genes for each factor between human and mouse reprogramming. Targets were defined as a gene whose TSS is closest to the peaks for each factor irrespective of binding distance. Because there were tens of thousands of O, S, K, and M peaks, about 40% to 70% of all genes could be assigned to O, S, K or M peak. We calculated the number of overlapping target genes among the four factors in the two species and found that a large fraction of genes was targeted by the four factors in both species (Fig 1b). Furthermore, among the 8,433 OSKM co-targeted genes in human and 6,867 co-targeted genes in mouse, 3,919 of them were shared significantly (p-value < 10^−16^, hypergeometric test), indicating a large fraction of OSKM co-targeted genes are conserved. Gene ontology enrichment analyses showed those shared co-targeted genes were enriched in the biological processes of regulation of transcription, in utero embryonic development and regulation of Wnt signaling pathway. This agrees with previous studies which showed that the Wnt signaling pathway modulated reprogramming efficiency when altered early in reprogramming (Ho, Papp, Hoffman, Merrill, & Plath, 2013). When only considering orthologous genes between human and mouse, we also found a large overlap of target genes for each reprogramming factor (Fig 1c). The hypergeometric test showed that the number of overlapping target genes was also significant for each of the four factors (p-value<10^−16^). Those results indicate that the factors tend to generally target the same genes in mouse and human fibroblasts. We also used another definition of target genes, requiring that the peak of each factor was within 20kb of the TSS and obtained a similar result (Supp Fig 7).

We next carried out *de novo* motif discovery in each factor’s binding regions (see Methods). The DNA binding motifs we identified for each reprogramming factor was similar between human and mouse (Fig 1d). However, we observed minor motif differences in Oct4, which terminated with A/T AA in mouse but A/G C/T AT in human, as well as in c-Myc, which terminated with C G/A TG in mouse but C/T G T/C G in human. Moreover, *de novo* motifs of the four factors were largely consistent with their canonical motifs (obtained from Jaspar database) (Mathelier et al., 2014), indicating DNA binding preferences of O, S, K, and M are largely conserved between human and mouse.

To further characterize OSKM binding, we identified all possible combinations of binding events. If summits of peaks from different reprogramming factors were within 100 bp of each other, we considered them to be “co-” binding events. If summits of peaks from one factor were at least 500 bp away from all other factors, we defined these as “solo” binding events. To gauge whether co-bound sites occurred more or less frequently than expected, we compared our counts to a synthetic null model for all possible combinations of factors (see Methods). We found that in both human and mouse, all co-binding events occurred more frequently than expected, whereas solo binding sites were observed less frequently than expected (Fig 1e). OSKM, OSK, OSM, OKM and SKM co-binding events were the most prevalent combinations in both human and mouse. Moreover, solo binding sites were more likely in human than in mouse and nearly all co-binding events (except KM and OKM) were more prevalent in mouse. Regardless of the differences, these results indicate that O, S, K, and M tend to bind together with similar combinatorial patterns in both human and mouse, suggesting that the factors often co-bind to exert their actions. Overall, we conclude that the general properties of O, S, K and M are similar, although there are some observable differences.

### Comparison of OSKM binding to the chromatin state of starting cells

Next, we sought to compare the chromatin state in the starting cells for OSKM binding sites at 48 hours between human and mouse. This enabled us to see how OSKM interacted with the initial chromatin states in fibroblasts. We analyzed H3K4me1, H3K4me3, H3K27me3, H3K27ac and H3K36me3 histone marks of human fibroblasts from the Roadmap Epigenome Project (Roadmap Epigenomics Consortium et al., 2015) and of mouse fibroblasts (Chronis et al., 2017) to build a 15 chromatin state model using ChromHMM (Ernst & Kellis, 2012) with a concatenated human-mouse genome (see Methods). Based on the combinatorial probability of the five histone marks, we classified the mouse and human genomes into chromatin states such as active promoter and active enhancer. We chose a model with 15 chromatin states because these had a clearly distinct combination of histone marks and functional annotations based on prior expectations. The genomes of both human and mouse were segmented into non-overlapping 200 bp regions, and each bin associated with a specific chromatin state. Figure 2a shows the emission probabilities (signal enrichments) of each histone mark as well as the fractions of the genome (numbers in the brackets, human followed by mouse) that each chromatin state occupies in human and mouse fibroblasts. We noted primary differences between human and mouse chromatin states including the frequency of the two H3K9me3-containing chromatin states, weak repressed polycomb and quiescent chromatin state, where human fibroblasts had significantly more genomic regions annotated as ZNF/repeats and heterochromatin and less genomic regions annotated as the latter two states.

**Figure 2.**
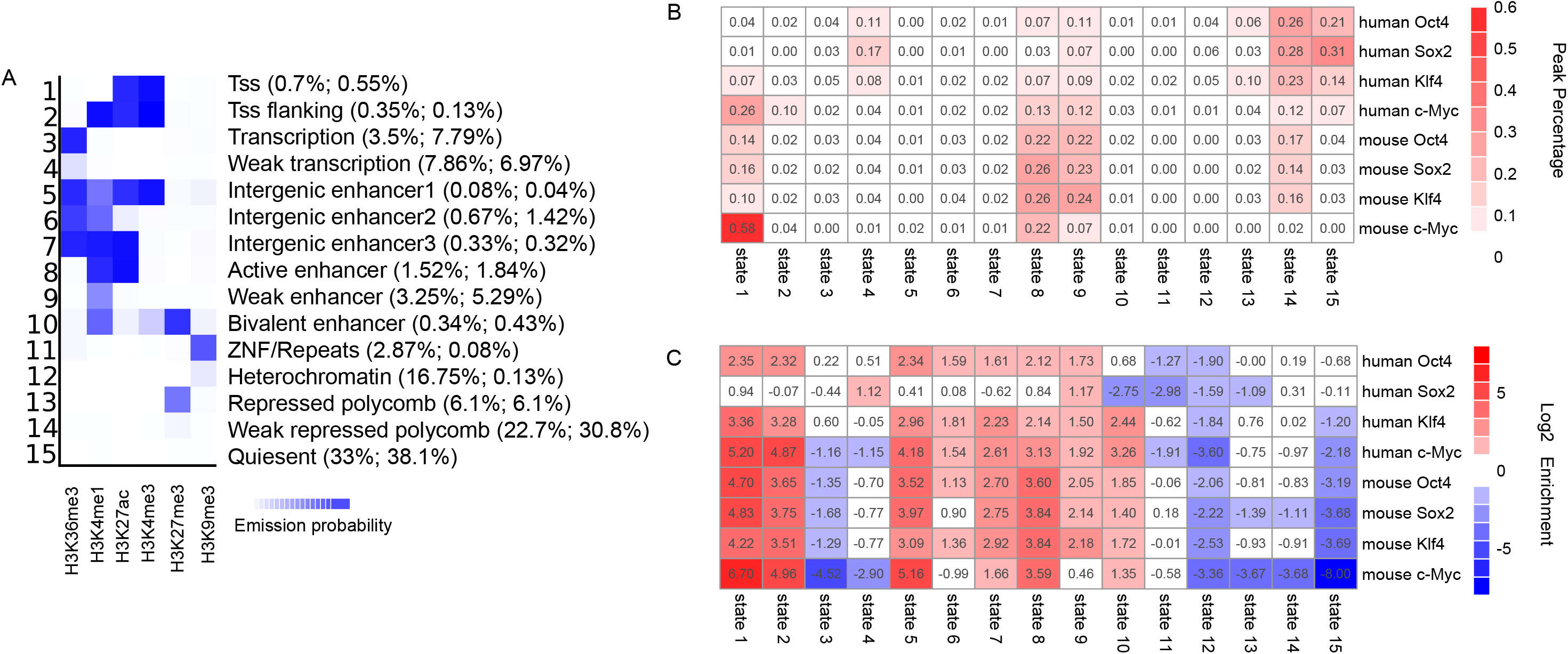
OSKM peaks target of the chromatin states in starting cells. A. Chromatin state model for concatenated human and mouse fibroblast cells based on five histone marks. The value in the heatmap represents the enrichment of that histone mark in that learned chromatin state. The values in the bracket represent the genomic percentage (human then followed by mouse) occupied by that chromatin state. B. Heatmap for percentages of OSKM peaks in each chromatin states from A. C. Heatmap for log2 enrichments between OSKM peaks percentages and chromatin state genomic percentages.

By intersecting OSKM peaks with chromatin states, we calculated the percentage of peaks within chromatin states in both human and mouse (Fig 2b). As a result, about 40~50% of human O, S, and K peaks, and 20% of human M peaks were within low signal regions (states 14 and 15). This chromatin analyses agrees with a direct assessment of the individual histone modification states targeted by OSKM, which showed that O, S, and K predominantly target unmarked chromatin sites (Soufi et al., 2012). By contrast, in mouse, the percentage of low signal regions targeted decreased to 20% for O, S, and K peaks and 2% for M peaks. In addition, about 40~50% of mouse O, S, and K peaks and 30% of M peaks were within enhancers, consistent with the finding that mouse OSK efficiently target enhancers active in fibroblasts early in mouse reprogramming (Chronis et al., 2017). However, for human O, S and K peaks, this number dropped to about 10%~25% and M peaks showed a similar number of 30%. After correction for the genome percentage annotated as different chromatin states, the human peaks were still more enriched in low signal regions and less enriched in enhancer regions. Those results reveal a distinct distribution of OSKM in chromatin states of low signals and enhancers between human and mouse. This binding preference may also suggest pluripotent genes in human are more difficult to induce and thus, human reprogramming will take longer than in mouse. In addition, consistent with the genomic distribution analysis (Fig 1a), mouse c-Myc was more often associated with promoters compared to human.

Additionally, when considering the genomic region captured by each chromatin state, we calculated the enrichment of OSKM peaks in each chromatin state by calculating the log2 ratio between peak percentage and chromatin state percentage (Fig 2c). This reveals OSKM binding preferences in the various chromatin states. As a result, we observed a strong preference of mouse OSKM targeting promoters and enhancers, whereas in human, this preference still held but the extent was decreased.

### OSKM binding events show limited conservation between human and mouse

To further compare OSKM occupancy in early mouse and human cell reprogramming, we mapped mouse peaks to the human genome based on synteny (see Methods). Mouse peaks were classified into three groups based on sequence conservation and binding conservation. Figure 3a shows a schematic illustration of the definition of the three groups: syntenic conserved peaks, syntenic unconserved peaks, and unsyntenic peaks. Syntenic conserved (SC) peaks had orthologous DNA sequences as well as binding events in both organisms. Syntenic unconserved (SU) peaks only had orthologous DNA sequences but no binding event detected in human. Unsyntenic (UN) peaks did not have orthologous DNA sequences between organisms and therefore could not be mapped between human and mouse.

**Figure 3.**
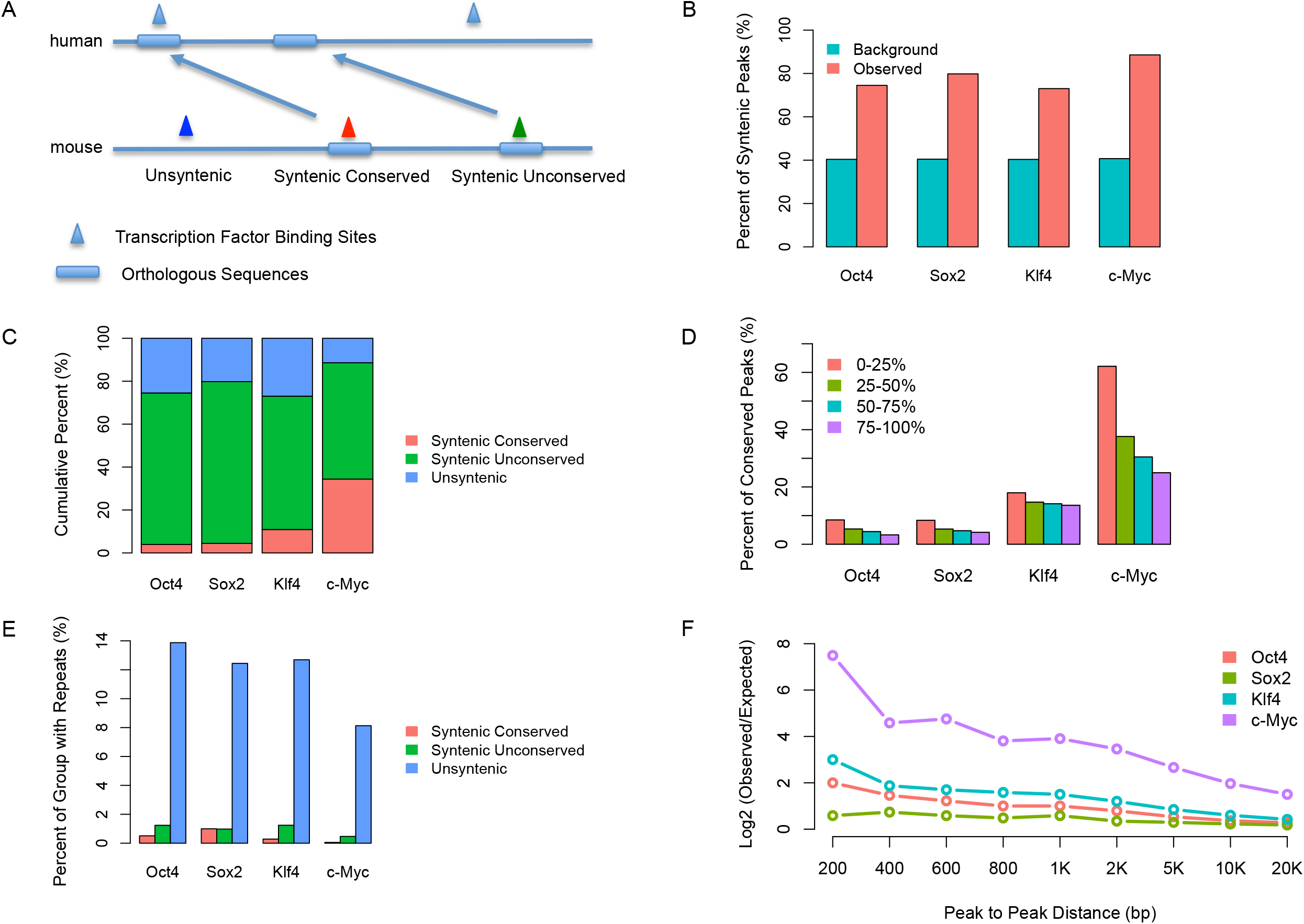
Map OSKM binding between human and mouse. A. Schematic illustration of the three different groups of peaks, i.e. Syntenic Conserved (SC) binding group, Syntenic Unconserved (SU) binding group and UNsyntenic (UN) binding group. B. Percentage of mouse OSKM peaks that can be mapped to human. The background is calculated by the simulation of peaks that have the same size and same number as the real peaks, and are allowed to map anywhere on the genome. C. Fractional constitutions of SC, SU and UN peaks for each factor. D. Percentage of SC binding events with respect to all syntenic binding events. For each factor, syntenic peaks are classified into four groups based on their peak enrichments of −log10(q-value). 0–25% are the top 25 percent of peaks while 75–100 are the bottom 25 percent of peaks. E. Percentage of the three groups of peaks that contain repeat sequences. F. Log2 fold enrichment of distances between human syntenic peaks in mouse and mouse peaks compared to random background.

We found that about 74, 80, 73 and 89 percent of mouse O, S, K, and M peaks, respectively, were syntenic with human, while the background ratio for the entire genome was about 40 percent (Fig 3b), indicating that elements bound by OSKM show much higher sequence conservation rates than the rest of the genome, consistent with OSKM bind to cis-regulatory events such as enhancers and promoters. However, for each reprogramming factor, we found that syntenic conserved peaks only represented a small fraction of peaks (Fig 3c). Specifically, 4%, 4.5%, 10.9% and 34.4% of mouse O, S, K, and M peaks, respectively, were syntenic conserved. O, S, and K, which mostly bind to enhancer regions in mouse (Fig 2b) (Chronis et al., 2017), had a lower fraction of conserved peaks compared to M, which mostly binds to promoter regions in early mouse cell reprogramming (Fig 2b) (Chronis et al., 2017). We then asked whether the limited degree of conservation between mouse and human binding events could be solely explained by random background binding events between human and mouse. To address this we simulated both human and mouse background peaks (same number and length with the observed ones), then calculated the conservation rate and repeated the simulation 1,000 times. The simulation result showed a conservation rate for OSKM background peaks of approximately 1%, implying that although the fraction of conserved binding was relatively small, conserved binding events still occurred at a higher rate than expected by chance. Lastly, we also mapped mouse pMX peaks (individual retroviral based system) to human peaks. Consistent with the comparison between polycistronic peaks in mouse and lentiviral peaks in human, our result showed that there was a limited fraction of syntenic conserved peaks for Oct4, Sox2 and Klf4 (Supp Fig 8). This result also indicates that the divergence of binding between human and mouse is not affected by using different overexpression systems.

In a previous study, Cheng et al. showed that the degree of binding conservation varied markedly, from several percent to about 60 percent, between human and mouse among different transcription factors (TFs) (Cheng et al., 2014). In addition, promoter bound TF binding sites showed higher conservation rates than enhancer sites. Moreover, this trend held after adjusting the sequence conservation differences between promoters and enhancers, indicating that the TF binding sites in promoter regions are indeed more conserved than those in enhancer regions (Cheng et al., 2014). In another study, Schmidt et al. reported a 10 to 22 percent binding conservation rate between two of five mammals for liver-specific transcription factors (Schmidt et al., 2010). In early reprogramming, we observed a low conservation rate for O, S, and K and a medium conservation rate for M, indicating the significance of binding divergence in early reprogramming system between human and mouse fibroblasts.

We next investigated whether peak binding strength (based on peak calling q-values) had an impact on conservation. We classified all mouse peaks into four groups based on their −log10 q-values (Fig 3d). For each reprogramming factor, we observed a clear trend where the strongest peaks (top 25%) had a higher percentage of syntenic conserved binding events compared to other three groups. This result suggests peak binding strength indeed is positively correlated with peak conservation rates and stronger peaks tend to be more conserved.

By analyzing the presence of repeat sequences within the three groups of peaks (see Methods) (SC, SU, and UN), we found that the unsyntenic peaks had a much higher percentage of repeat sequences compared to the other two groups, and, except for Sox2, syntenic conserved binding sites contained the fewest repeats (Fig 3e). Moreover, compared to peaks in syntenic regions, peaks in unsyntenic regions were more often associated with long terminal repeats (LTR) and short interspersed nuclear elements (SINE) and less often with simple repeats in the mouse genome (Supp Fig 9). These results are consistent with previous findings which showed that transposable elements are enriched in species-specific sequences and have rewired the transcriptional network during evolution (Kunarso et al., 2010; Sundaram et al., 2014).

The analyses described above were carried out by mapping mouse OSKM peaks to the human genome, but we also performed the inverse analysis by mapping human OSKM peaks to the mouse genome (Supp Fig 10a). Approximately 60 percent of human peaks occurred in genomic regions syntenic with the mouse. The lower syntenic rate of human peaks mapping to the mouse genome compared with mouse peaks mapping to the human genome correlated with a higher proportion of repeats in human peak sequences (Fig 9). Among human OSKM peaks in syntenic regions, those also found in the mouse (syntenic conserved) constituted a small proportion as seen in the reverse mapping of mouse OSKM peaks to the human data (Supp Fig 10b). Interestingly, syntenic and unsyntenic human OSKM peaks showed a more similar distribution of certain types of repeats compared to mouse peaks (Supp Fig 11, Supp Fig 12).

We also investigated how human syntenic peaks and all peaks of mouse were distributed relative to each other. We first calculated the distances between human syntenic peak summits and mouse peak summits. We then categorized the distances into several groups of genomic ranges, i.e. within 200bp, 400bp, 600bp, 800bp etc. Lastly, to compare the observed distance distribution with simulated background, we calculated the background distance distribution, where the mouse peaks were shuffled and the human syntenic peaks were kept fixed. The result suggests that observed human syntenic peaks are indeed closer to observed mouse peaks than expected by chance (Fig 3f). Moreover, there was a clear trend showing that the log2 ratio between observed and simulated peaks declined with increased distance. Among the four factors, c-Myc showed the most dramatic trend. This is consistent with the fact that c-Myc is the most conserved factor compared to the other three.

### Syntenic conserved peaks are associated with different genomic features compared with unconserved peaks

Since we observed that only a small fraction of syntenic peaks had conserved binding early in reprogramming in human and mouse cells, we sought to identify properties that distinguish conserved peaks from the others. We observed that syntenic conserved peaks had significantly higher ChIP enrichment (−log10 q-value) than the other two groups (Fig 4a), indicating the SC peaks tend to be bound more strongly. We then used the GREAT tool (McLean et al., 2010) to perform gene ontology enrichment analysis for the mouse SC, SU, and UN peaks, with all peaks as background (Supp Fig 13). For SC peaks of OSM, we found their target genes were enriched for fat pad, adipose tissue, and adrenal gland development. Surprisingly, for SU peaks no enriched gene ontology terms for any of the four factors were detected. UN peaks of OSKM were strongly enriched in immunity-related gene ontologies. These results suggest that the target genes of the three groups of peaks might be associated with distinct functions. When comparing the genomic locations of mouse SC peaks to all peaks with respect to the distance to the TSSs, we found that SC O, K, and M peaks more often occurred within the proximal TSS regions, while Sox2 was slightly more often within the distal TSS regions (Fig 4b).

**Figure 4.**
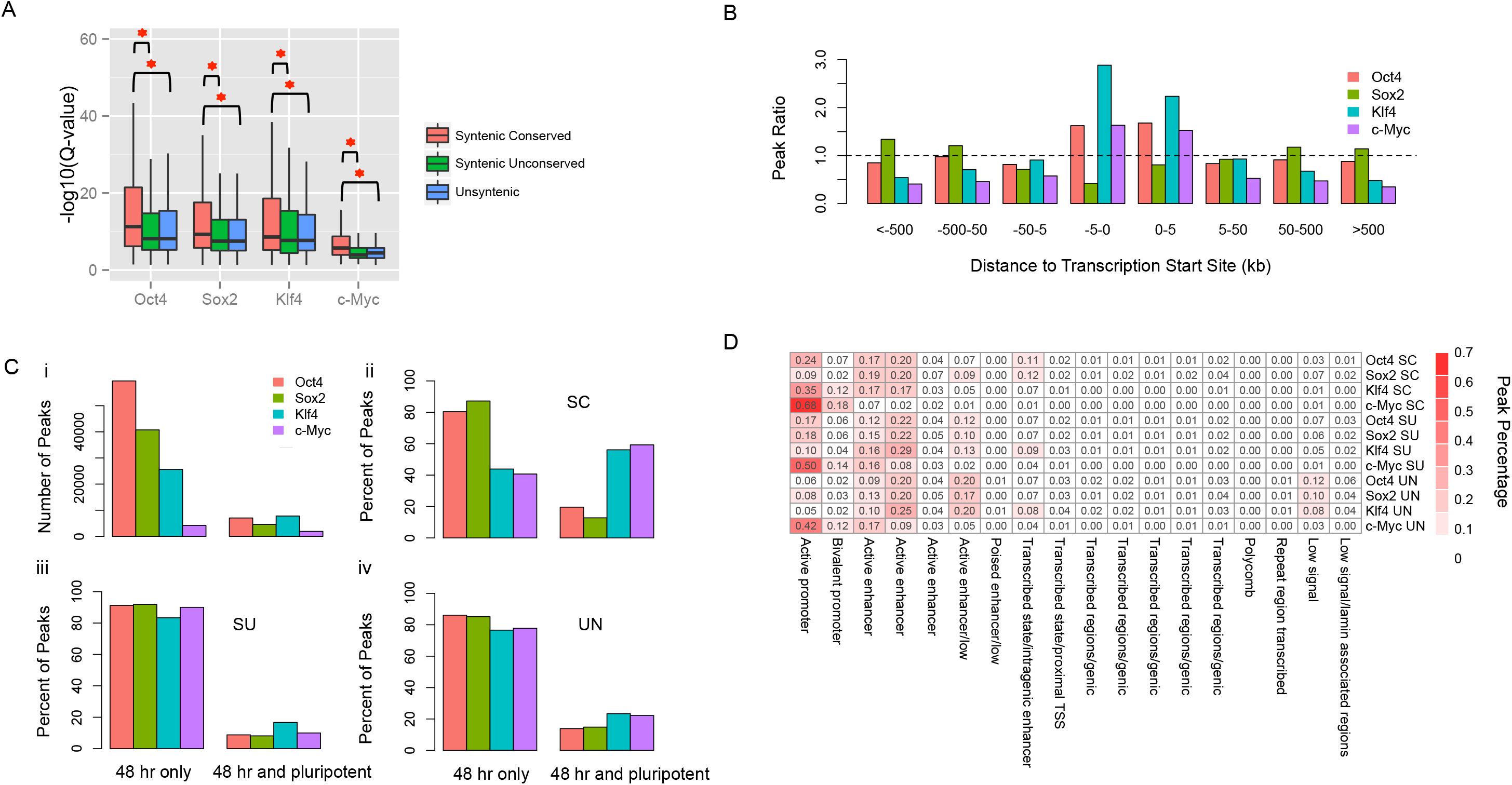
Comparisons of syntenic conserved peaks with syntenic unconserved peaks and unsyntenic peaks. A. Box plot of peak calling q-values for the SC, SU and UN groups of peaks. B. Fold enrichment of positional distribution between SC peaks and all peaks around Transcription Start Sites. C. Percentage of SC, SU and UN peaks with consecutive bindings. 48 hr only represents the peaks that only bound in 48 hours of reprogramming, while 48 hr and pluripotent represents the peaks that are also bound in the reprogramming final stage. i represents the number of the two group of peaks. ii–iv represents the percentage of SC, SU and UN peaks that are either 48 hr only bound or 48 hr and pluripotent bound. D. Heatmap for percentages of mouse SC, SU and UN peaks in the mouse 18 chromatin states.

We also compared binding of mouse OSKM at 48 hours with that in the pluripotent state, to define those mouse OSKM binding events that were bound both early in reprogramming and in the pluripotent state (based on mouse embryonic stem cell ChIP-seq data) versus those that only occur at 48 hours but not in pluripotent cells (Fig 4c,i) (Chronis et al., 2017). In our previous study, we described that many of these persistent binding events for OSKM were enriched in promoters and OSK were also highly enriched in pluripotency enhancers (Chronis et al., 2017). We calculated the percentage of SC peaks that were bound only early in reprogramming or persist throughout reprogramming. We found that compared with SU and UN peaks, mouse SC peaks of OKM at 48 hours had a higher fraction of persistent binding events (Fig 4c,ii–iv). Specifically, for Oct4, the percent of persistent bound events was 20, 9 and 14 for SC, SU and UN respectively. For Klf4, this percent was 56, 17 and 23, and for c-Myc, this percent was 59, 10 and 22. This result indicates that conserved binding events, especially for K and M, tend to be maintained during reprogramming and are therefore likely to be more functionally important than unconserved ones.

We next asked whether SC, SU and UN peaks had distinct patterns of chromatin states in mouse at 48 hours. A mouse 18 chromatin state model was generated with nine histone marks and described in our previous paper (Supp Fig 14) (Chronis et al., 2017). We therefore calculated the percentage of peaks within each chromatin state (Fig 4d). As a result, we found that SC peaks preferentially tended to occur within certain chromatin states compared to SU and UN peaks. Specifically, SC peaks of O, K and M had higher percentages within active promoters, bivalent promoters and certain groups of enhancers. By contrast, UN peaks of O, S and K had higher percentages within low signal regions. Those results indicate that different groups of peaks are likely to associate with different chromatin states.

To further investigate the chromatin states of syntenic peaks, we performed another comparison from a human-mouse transition perspective. We assigned each syntenic peak to the chromatin state in the concatenated human and mouse genome (Fig 2a) and compared the chromatin assignment of each SC peak between mouse and human (see Methods) (Fig 5a). The color in the heatmap reflects the percentage of SC peaks within that transition in chromatin state between the mouse and human syntenic genome. For example, the top left square in the heatmap is the transition from human TSS regions (state 1) to mouse TSS regions (state 1) and the bottom right is the transition from human quiescent regions (state 15) to mouse quiescent regions (state 15) (i.e. no changes in chromatin state), and any deviation from the diagonal represents a change in chromatin state. For SC peaks of O, S, and K, the most frequent transitions corresponded to human promoter to mouse promoter, human enhancer to mouse promoter, human enhancer to mouse enhancer, and human enhancer to mouse quiescent regions. By contrast, the majority of frequent transitions for c-Myc involved promoter to promoter states. We also asked whether the chromatin state transition patterns were different for unconserved peaks. When comparing the transition profiles between SC and SU peaks (Fig 5b), we found an enrichment in human promoter to mouse promoter, human enhancer to mouse promoter and human enhancer to mouse enhancer transitions, indicating that SC peaks are more often associated with certain regulatory sites in both species than SU peaks.

**Figure 5.**
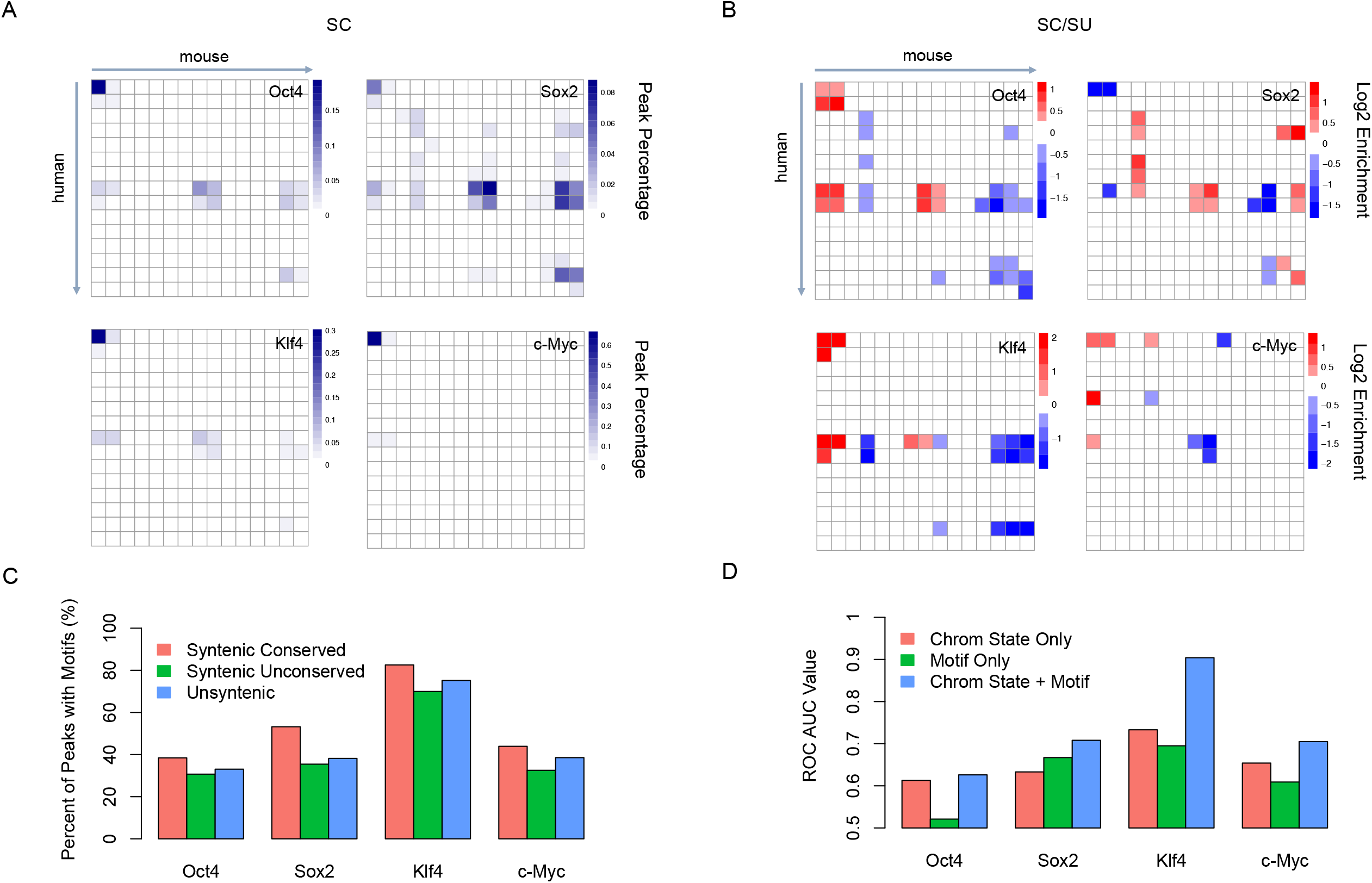
Chromatin state transitions, motif usages and their contributions in the maintaining of syntenic conserved peaks. A. Chromatin state transitions of syntenic conserved peaks between human and mouse. The top left is state 1 of human to state 1 of mouse. The value in the heatmap represents the fraction of the number of syntenic conserved peaks in that square divided by the total number of all syntenic conserved peaks. B. Chromatin state transitions of the log2 ratio between syntenic conserved peaks versus syntenic unconserved peaks. The value in the heatmap represents the log2 ratio between the fraction of syntenic conserved peaks and the fraction of syntenic unconserved peaks in that square. C. Percentage of SC, SU and UN peaks that have canonical motifs. D. ROC AUC of a classifier to predict syntenic based on motif occurrences and chromatin state transitions.

Another factor that may help maintain the conservation of peaks is the occurrence of binding motifs. Although we observed that SC peaks were preferentially found within promoters and enhancers, it was not clear whether motifs help maintain the conservation of peaks between mouse and human. To shed light on this question, we computed the motif frequency in each group of peaks (see Methods) (Fig 5c). We reasoned that if the conservation of peaks was strongly influenced by the presence of binding motifs between mouse and human, then SC peaks should have a different fraction of motifs compared to the other two groups. For Sox2, 53% of SC binding events had identifiable motifs within their peaks, compared to approximately 35% of SU and UN. However, for the other three factors, SC peaks contained more motifs but the differences among the three types of peaks were smaller, indicating the limited impact of sequence motifs in the determination of binding conservation.

### Using transitions of regulatory motifs and chromatin states as predictors of conserved binding

To quantitatively assess the extent to which SC peaks are determined by motifs or chromatin states, we built a naïve Bayesian classifier to evaluate the prediction power for classifying syntenic peaks into the SC and SU groups (see Methods). This model was trained using different sources of information: motif only, chromatin state only, and the two combined. Area under the curve (AUC) values of receiver operator curves (ROC) were used to estimate the prediction power (Fig 5d). We found that except for Sox2, the chromatin state only model outperformed the motif only model. Moreover, when combining information from both motif and chromatin states, the AUC for O, S, K, and M were 0.63, 0.71, 0.90, and 0.71 respectively. Klf4 showed a strikingly high prediction power due to its strong motif preference in syntenic regions between human and mouse and its strong chromatin state preference for specific chromatin state transitions. Although the models for O, S, and M only predicted a fraction of conserved sites, these results demonstrate that conserved peaks are indeed associated with syntenic regions that contain strong motif sequences and preferred chromatin state transitions between mouse and human.

## Discussion

In this study, we systematically compared binding patterns of the four reprogramming factors OSKM between human and mouse at an early time point of reprogramming to the iPSC state. When analyzed in each genome separately, OSKM binding sites in human and mouse shared similar features: OSK tend to bind distal TSS regions, OSKM tend to target similar genes, have similar DNA binding motifs, and show similar combinatorial binding patterns among the reprogramming factors. This suggests that molecular properties of these factors are conserved between human and mouse. However, differences emerged when we investigated the chromatin state of target sites: OSKM targeted far more closed (low signal state) chromatin states in human cells than in mouse. Importantly, when we compared the binding sites across syntenic regions, we found that there was only a small percentage of sites that were bound in both genomes (i.e. syntenic conserved, SC). Altogether, our results suggest that the initial OSKM binding sites are largely distinct in these two species, even though the phenotypic consequences of these interactions ultimately lead to similar cell types.

We also observed that most early binding events do not persist in the later stages of iPSCs reprogramming (Chronis et al., 2017). However, we found that binding events that were conserved between mouse and human tended to persist more often throughout the reprogramming process compared to unconserved sites. Conserved binding sites also tended to have a higher proportion of conserved cis-regulatory elements associated with each factor. We also showed that binding sites were more likely to be conserved if the mouse and human chromatin states were similar and the motifs were conserved.

We recognize that there are certain limitations to our analysis. One is that human and mouse reprogramming was performed using slightly different experimental protocols. An inducible polycistronic cassette including all four reprogramming factors was used in mouse fibroblasts, ensuring homogeneous expression and stoichiometry across the cell population at 48 hours; whereas four separate lentiviral constructs were used in human, each expressing one factor. However, as we have shown by comparing mouse polycistronic to individual cassettes, these different overexpression methods lead to very similar binding peaks. Also, it is possible that at 48 hours, human and mouse cells might not be in the same reprogramming stages due to their different reprogramming kinetics. However, the time point we used corresponds to early events in the time series of both species, and should, therefore, identify the first interactions of these factors with chromatin. Moreover, we compared mouse embryonic fibroblasts and human fetal foreskin fibroblasts as starting cells of reprogramming. However, we believe that the epigenome changes from embryonic to fetal stages of fibroblasts are unlikely to have a dramatic effect on OSKM binding patterns. As a result, our conclusions drawn from the comparison of these two species should not be significantly affected by the differences in the experimental details of the human and mouse systems.

In conclusion, we have shown that while some general properties of OSKM binding are conserved between mouse and human, the detailed transcriptional network is vastly reorganized. A subset of the binding events are syntenic between the two species and this study has allowed us to identify these. We do not know if they represent key events that are distinct from the large fraction of other binding sites that are not conserved. However, several lines of evidence that we have presented, such as the fact that these sites tend to persist throughout the reprogramming process, do suggest that these may play a more significant role in reprogramming than the typical unconserved site. Nonetheless, the overall picture that emerges is that the OSKM regulatory networks have significantly diverged between the two species, and while the general properties of these networks are similar, the specific binding sites are generally distinct. This observation may suggest that reprogramming to pluripotency may be driven by global regulatory changes in cells that do not depend critically on a small set of specific interactions.

## Methods

### Cell culture and reprogramming

In the human reprogramming system, BJ fibroblasts were obtained from ATCC (CRL-2522) at passage 6 and cultured in the ATCC-formulated Eagle’s Minimum Essential Medium supplemented with 10% fetal bovine serum at 37 C and 5% CO2. The human H1-ES line (Thomson et al., 1998) and our derived hFib-iPS cell lines were maintained as described (Lerou et al., 2008). More information about experimental details can be found in the supplementary documents of Soufi et al. 2012 (Soufi et al., 2012). In the mouse reprogramming system, mouse embryonic fibroblasts carrying a polycistronic, dox-inducible OSKM cassette in the Col1A locus and a heterozygous M2rtTA allele in the R26 locus, were grown in standard mouse ESC media containing knockout-DMEM, 15% fetal bovine serum, recombinant leukemia inhibitory factor (Lif), b-mercaptoethanol, 1x penicillin/streptomycin, L-glutamine, and non-essential amino acids. Repogramming was induced by the addition of 2ug/ml doxycycline. We generated mouse iPS cell lines as described (Hockemeyer et al., 2008; Park et al., 2008). Briefly, BJ cells at passage 10 were infected with lentiviruses encoding for dox-inducible Oct4, Sox2, Klf4, and c-Myc, along with lentiviruses expressing rtTA2M2 in the presence of 4.5 mg/ml polybrene. Additional experimental details can be found in the supplementary documents of Chronis et al. 2017 (Chronis et al., 2017).

### Mapping and Peak Calling

The human OSKM ChIP-Seq datasets were downloaded from GEO with accession number of GSE36570, while mouse OSKM ChIP-Seq datasets were downloaded from GEO with accession number of GSE90895. Bowtie was used to map ChIP-Seq reads of both human and mouse to their respective genomes allowing two mismatches and keeping only uniquely mapped reads for further analysis (Langmead, Trapnell, Pop, & Salzberg, 2009). MACS2 2.1.0 was used to identify ChIP-Seq peaks with a q-value cutoff of 0.05 (Zhang et al., 2008).

### Motif finding and motif occurrences within peaks

MEME-ChIP was used to perform de novo motif finding for OSKM binding peaks (Machanick & Bailey, 2011). To identify the strongest motifs, the identified summits of peaks were ranked based on their enrichments and the top 10,000 summits, along with their surrounding 200bp, were used as the input regions. The enriched motifs were identified using the DREME algorithm in the MEME-ChIP software. Starting with the most significant motif for each factor, we then used the Position Weight Matrix of this motif to scan for peaks, and determined the peaks associated with this motif using a p-value cutoff of 0.001.

### Combinatorial binding and solo binding

To identify combinatorial binding regions where multiple factors bind, peaks were merged if their summits were within 100 bp of each other. Then these different combinations of binding sites were broken down into their different combinations of factors. To identify solo binding regions where only one factor bound, we required that its summit be at least 500 bp away from all other factors. Note that this method is more stringent than that used by Soufi et al. (Soufi et al., 2012); the latter considered solo binding events as simply not falling within 100 bp of the peak center. Here, to estimate the background rates of combinatorial binding, the peaks of OSKM were first randomly shuffled in the genome (using the bedtools shuffle function) (Quinlan & Hall, 2010). Secondly, the expected number of combinatorial binding events was re-calculated based on these shuffled peaks. Lastly, we compared the number of observed binding events versus the number of expected binding events for all possible combinations of factors.

### RNA-Seq samples and analysis

The human fibroblasts and 48 hours of reprogramming cells for RNA-Seq were cultured and generated in the same condition with the samples for OSKM ChIP-Seq analyzed in this study. The total mRNA was then extracted and sequenced. The experimental details could be found within method section in Tong et al. (Tong et al., 2016). The RNA-Seq samples for mouse fibroblasts cells were obtained from Chronis et al. (Chronis et al., 2017). The raw sequencing reads of both human and mouse were then mapped back to their corresponding genome using Tophat (Trapnell, Pachter, & Salzberg, 2009). After this, HTSeq software was used to calculate the number of reads within each gene for both human and mouse (Anders, Pyl, & Huber, 2015). Finally, the DESeq2 software was used to perform the differential expressed gene analysis with a q-value cutoff of 0.05 (Love et al., 2014).

### Mapping sequences between human and mouse

To map OSKM binding sites between human and mouse, the liftOver algorithm from the UCSC Genome Browser was used with a cutoff of 0.5. The LiftOver algorithm uses an alignment chain file to map genomic coordinates between different versions of assemblies, or different species. The algorithm searches for regions where the input sequences are in the same block with the converted assemblies or species. The cutoff of 0.5 requires that the mapped sequences share at least half of exactly same DNA sequences with the converted species. This cutoff is consistent with modENCODE project paper which compares transcription factor binding sites between human and mouse (Cheng et al., 2014). To confirm the reliability of our results, we also used another method named bnMapper and got very similar results (Denas et al., 2015).

### Peaks associated with repeat sequences

Repeat sequences were downloaded from the RepeatMasker database. We extracted the genomic coordinates for the major repeat families including DNA (DNA transposon elements), LINE (Long interspersed nuclear elements), LTR (Long terminal repeats), Retroposon (Transposons via RNA intermediates), Satellite (Satellite DNA which belongs to tandem repeats), Simple (Simple repeats) and SINE (Short interspersed nuclear elements). A peak was considered to be associated with a repeat sequence if the genomic coordinate of this repeat was within this peak.

### Chromatin states for concatenated human and mouse genomes

For mouse histone marks, we used the datasets for mouse fibroblast cells from our previously published paper (Chronis et al., 2017). For human histone marks, we used the datasets of IMR90 fibroblast cell line downloaded from RoadMap Epigenomics Project (Roadmap Epigenomics Consortium et al., 2015). To learn the joint chromatin state for human and mouse, a pseudo chromosome size table was constructed by concatenating human and mouse genomes. Then the model was trained with the human fibroblast and mouse embryonic fibroblast histone data, producing a common set of emission probabilities. We then generated a 15 chromatin state model based on the combinatorial patterns of five histone marks, i.e, H3K4me1, H3K4me3, H3K27me3, H3K27ac and H3K36me3.

### Chromatin state transitions between human and mouse

Each syntenic conserved peak in mouse and its corresponding orthologous peak in human was assigned a chromatin state as described above. We then calculated the number of peaks within each possible chromatin state transition. This leads to the generation of a 15 × 15 chromatin state transition matrix. For example, the top left of the matrix represents the fraction of syntenic peaks with state 1 of human and state 1 of mouse. We also performed the same calculation for syntenic unconserved peaks between human and mouse. To compare to relative enrichment of chromatin state transitions, the log2 ratio between the syntenic conserved and syntenic unconserved matrices was calculated.

### Classification model

We built a Naïve Bayes model to classify syntenic peaks into a syntenic conserved and syntenic unconserved group, based on their chromatin state transition (see above) and motif occurrences transitions. The motif occurrence transition matrix was a 2 × 2 matrix that represents the frequency of motif occurrences for syntenic peaks between human and mouse. Log odds ratios were then calculated between syntenic conserved group and syntenic unconserved groups for both chromatin state transition and motif occurrence transition matrices. As a result, each peak was assigned two values: one was the chromatin state transition log odds ratio matrix to represent the chromatin state model, and another was the motif occurrences transition log odds ratio matrix to represent the motif model. The two values were added to represent both the chromatin state and motif occurrence model. Syntenic peaks were then ranked based on log odds ratio values from either the chromatin state transition matrix or motif occurrences transition matrix, or their sum. A syntenic conserved peak was labeled as 1 and a syntenic unconserved peak is labeled as 0. Lastly, the Area under the curve (AUC) values of the receiver operator curves (ROC) were calculated to represent the model performance for classifying syntenic peaks into 1 or 0 given the chromatin state transitions or motif occurrences transitions.

### Author contribution

M.P., K.P., K.Z and K.F. designed the research plan; K.F. performed the data analysis; C.C, A.S. and M.E. performed the experiments; K.F. and M.P wrote the manuscript; K.Z and K.P supervised the study; K.Z., K.P., C.C. and A.S. contributed to writing; G.B and S.S contributed to discussion.

## Acknowledgment

This work was supported by P01 grant from NIH (grant number: 5P01GM099134).

## References

Anders, S., Pyl, P. T., & Huber, W. (2015). HTSeq--a Python framework to work with high-throughput sequencing data. Bioinformatics (Oxford, England), 31(2), 166–169. http://doi.org/10.1093/bioinformatics/btu638

Cheng, Y., Ma, Z., Kim, B.-H., Wu, W., Cayting, P., Boyle, A. P., et al. (2014). Principles of regulatory information conservation between mouse and human. Nature, 515(7527), 371–375. http://doi.org/10.1038/nature13985

Chronis, C., Fiziev, P., Papp, B., Butz, S., Bonora, G., Sabri, S., et al. (2017). Cooperative Binding of Transcription Factors Orchestrates Reprogramming. Cell, 168(3), 442–459.e20. http://doi.org/10.1016/j.cell.2016.12.016

Denas, O., Sandstrom, R., Cheng, Y., Beal, K., Herrero, J., Hardison, R. C., & Taylor, J. (2015). Genome-wide comparative analysis reveals human-mouse regulatory landscape and evolution. BMC Genomics, 16(1), 87. http://doi.org/10.1186/s12864-015-1245-6

Ernst, J., & Kellis, M. (2012). ChromHMM: automating chromatin-state discovery and characterization. Nature Methods, 9(3), 215–216. http://doi.org/10.1038/nmeth.1906

Hirschi, K. K., Li, S., & Roy, K. (2014). Induced pluripotent stem cells for regenerative medicine. Annual Review of Biomedical Engineering, 16(1), 277–294. http://doi.org/10.1146/annurev-bioeng-071813-105108

Ho, R., Papp, B., Hoffman, J. A., Merrill, B. J., & Plath, K. (2013). Stage-specific regulation of reprogramming to induced pluripotent stem cells by Wnt signaling and T cell factor proteins. Cell Reports, 3(6), 2113–2126. http://doi.org/10.1016/j.celrep.2013.05.015

Hockemeyer, D., Soldner, F., Cook, E. G., Gao, Q., Mitalipova, M., & Jaenisch, R. (2008). A drug-inducible system for direct reprogramming of human somatic cells to pluripotency. Cell Stem Cell, 3(3), 346–353. http://doi.org/10.1016/j.stem.2008.08.014

Koche, R. P., Smith, Z. D., Adli, M., Gu, H., Ku, M., Gnirke, A., et al. (2011). Reprogramming factor expression initiates widespread targeted chromatin remodeling. Cell Stem Cell, 8(1), 96–105. http://doi.org/10.1016/j.stem.2010.12.001

Kunarso, G., Chia, N.-Y., Jeyakani, J., Hwang, C., Lu, X., Chan, Y.-S., et al. (2010). Transposable elements have rewired the core regulatory network of human embryonic stem cells. Nature Genetics, 42(7), 631–634. http://doi.org/10.1038/ng.600

Langmead, B., Trapnell, C., Pop, M., & Salzberg, S. L. (2009). Ultrafast and memory-efficient alignment of short DNA sequences to the human genome. Genome Biology, 10(3), R25. http://doi.org/10.1186/gb-2009-10-3-r25

Lerou, P. H., Yabuuchi, A., Huo, H., Miller, J. D., Boyer, L. F., Schlaeger, T. M., & Daley, G. Q. (2008). Derivation and maintenance of human embryonic stem cells from poor-quality in vitro fertilization embryos. Nature Protocols, 3(5), 923–933. http://doi.org/10.1038/nprot.2008.60

Love, M. I., Huber, W., & Anders, S. (2014). Moderated estimation of fold change and dispersion for RNA-seq data with DESeq2. Genome Biology, 15(12), 550. http://doi.org/10.1186/s13059-014-0550-8

Machanick, P., & Bailey, T. L. (2011). MEME-ChIP: motif analysis of large DNA datasets. Bioinformatics (Oxford, England), 27(12), 1696–1697. http://doi.org/10.1093/bioinformatics/btr189

Malik, N., & Rao, M. S. (2013). A review of the methods for human iPSC derivation. Methods in Molecular Biology (Clifton, N.J.), 997(Chapter 3), 23–33. http://doi.org/10.1007/978-1-62703-348-0_3

Mathelier, A., Zhao, X., Zhang, A. W., Parcy, F., Worsley-Hunt, R., Arenillas, D. J., et al. (2014). JASPAR 2014: an extensively expanded and updated open-access database of transcription factor binding profiles. Nucleic Acids Research, 42(Database issue), D142–7. http://doi.org/10.1093/nar/gkt997

McLean, C. Y., Bristor, D., Hiller, M., Clarke, S. L., Schaar, B. T., Lowe, C. B., et al. (2010). GREAT improves functional interpretation of cis-regulatory regions. Nature Biotechnology, 28(5), 495–501. http://doi.org/10.1038/nbt.1630

Nichols, J., & Smith, A. (2009). Naive and primed pluripotent states. Cell Stem Cell, 4(6), 487–492. http://doi.org/10.1016/j.stem.2009.05.015

Ohnuki, M., Tanabe, K., Sutou, K., Teramoto, I., Sawamura, Y., Narita, M., et al. (2014). Dynamic regulation of human endogenous retroviruses mediates factor-induced reprogramming and differentiation potential. Proceedings of the National Academy of Sciences of the United States of America, 111(34), 12426–12431. http://doi.org/10.1073/pnas.1413299111

Park, I.-H., Zhao, R., West, J. A., Yabuuchi, A., Huo, H., Ince, T. A., et al. (2008). Reprogramming of human somatic cells to pluripotency with defined factors. Nature, 451(7175), 141–146. http://doi.org/10.1038/nature06534

Polo, J. M., Anderssen, E., Walsh, R. M., Schwarz, B. A., Nefzger, C. M., Lim, S. M., et al. (2012). A molecular roadmap of reprogramming somatic cells into iPS cells. Cell, 151(7), 1617–1632. http://doi.org/10.1016/j.cell.2012.11.039

Quinlan, A. R., & Hall, I. M. (2010). BEDTools: a flexible suite of utilities for comparing genomic features. Bioinformatics (Oxford, England), 26(6), 841–842. http://doi.org/10.1093/bioinformatics/btq033

Roadmap Epigenomics Consortium, Kundaje, A., Meuleman, W., Ernst, J., Bilenky, M., Yen, A., et al. (2015). Integrative analysis of 111 reference human epigenomes. Nature, 518(7539), 317–330. http://doi.org/10.1038/nature14248

Schmidt, D., Wilson, M. D., Ballester, B., Schwalie, P. C., Brown, G. D., Marshall, A., et al. (2010). Five-vertebrate ChIP-seq reveals the evolutionary dynamics of transcription factor binding. Science (New York, N.Y.), 328(5981), 1036–1040. http://doi.org/10.1126/science.1186176

Singh, V. K., Kalsan, M., Kumar, N., Saini, A., & Chandra, R. (2015). Induced pluripotent stem cells: applications in regenerative medicine, disease modeling, and drug discovery. Frontiers in Cell and Developmental Biology, 3, 2. http://doi.org/10.3389/fcell.2015.00002

Soufi, A., Donahue, G., & Zaret, K. S. (2012). Facilitators and impediments of the pluripotency reprogramming factors’ initial engagement with the genome. Cell, 151(5), 994–1004. http://doi.org/10.1016/j.cell.2012.09.045

Soufi, A., Garcia, M. F., Jaroszewicz, A., Osman, N., Pellegrini, M., & Zaret, K. S. (2015). Pioneer transcription factors target partial DNA motifs on nucleosomes to initiate reprogramming. Cell, 161(3), 555–568. http://doi.org/10.1016/j.cell.2015.03.017

Sundaram, V., Cheng, Y., Ma, Z., Li, D., Xing, X., Edge, P., et al. (2014). Widespread contribution of transposable elements to the innovation of gene regulatory networks. Genome Research, 24(12), gr.168872.113-1976. http://doi.org/10.1101/gr.168872.113

Takahashi, K., & Yamanaka, S. (2006a). Induction of pluripotent stem cells from mouse embryonic and adult fibroblast cultures by defined factors. Cell, 126(4), 663–676. http://doi.org/10.1016/j.cell.2006.07.024

Takahashi, K., & Yamanaka, S. (2006b). Induction of pluripotent stem cells from mouse embryonic and adult fibroblast cultures by defined factors. Cell, 126(4), 663–676. http://doi.org/10.1016/j.cell.2006.07.024

Takahashi, K., & Yamanaka, S. (2016). A decade of transcription factor-mediated reprogramming to pluripotency. Nature Reviews. Molecular Cell Biology, 17(3), 183–193. http://doi.org/10.1038/nrm.2016.8

Takahashi, K., Tanabe, K., Ohnuki, M., Narita, M., Ichisaka, T., Tomoda, K., & Yamanaka, S. (2007). Induction of pluripotent stem cells from adult human fibroblasts by defined factors. Cell, 131(5), 861–872. http://doi.org/10.1016/j.cell.2007.11.019

Thomson, J. A., Itskovitz-Eldor, J., Shapiro, S. S., Waknitz, M. A., Swiergiel, J. J., Marshall, V. S., & Jones, J. M. (1998). Embryonic stem cell lines derived from human blastocysts. Science (New York, N.Y.), 282(5391), 1145–1147.

Tong, A.-J., Liu, X., Thomas, B. J., Lissner, M. M., Baker, M. R., Senagolage, M. D., et al. (2016). A Stringent Systems Approach Uncovers Gene-Specific Mechanisms Regulating Inflammation. Cell, 165(1), 165–179. http://doi.org/10.1016/j.cell.2016.01.020

Trapnell, C., Pachter, L., & Salzberg, S. L. (2009). TopHat: discovering splice junctions with RNA-Seq. Bioinformatics (Oxford, England), 25(9), 1105–1111. http://doi.org/10.1093/bioinformatics/btp120

Yamanaka, S. (2012). Induced pluripotent stem cells: past, present, and future. Cell Stem Cell, 10(6), 678–684. http://doi.org/10.1016/j.stem.2012.05.005

Yang, C.-S., Chang, K.-Y., & Rana, T. M. (2014). Genome-wide functional analysis reveals factors needed at the transition steps of induced reprogramming. Cell Reports, 8(2), 327–337. http://doi.org/10.1016/j.celrep.2014.07.002

Zhang, Y., Liu, T., Meyer, C. A., Eeckhoute, J., Johnson, D. S., Bernstein, B. E., et al. (2008). Model-based analysis of ChIP-Seq (MACS). Genome Biology, 9(9), R137. http://doi.org/10.1186/gb-2008-9-9-r137

